# The genomes of the *Macadamia* genus

**DOI:** 10.1101/2023.12.07.570730

**Authors:** Priyanka Sharma, Ardashir Kharabian Masouleh, Lena Constantin, Bruce Topp, Agnelo Furtado, Robert J. Henry

## Abstract

*Macadamia*, a genus native to Eastern Australia, comprises four species, *Macadamia integrifolia, M. tetraphylla, M. ternifolia,* and *M. jansenii*. Macadamia was recently domesticated largely from a limited gene pool of Hawaiian germplasm and has become a commercially significant nut crop. Disease susceptibility and climate adaptability challenges, highlight the need for use of a wider range of genetic resources for macadamia production. High quality haploid resolved genome assemblies were generated using HiFiasm to allow comparison of the genomes of the four species. Assembly sizes ranged from 735 Mb to 795 Mb and N50 from 53.7 Mb to 56 Mb, indicating high assembly continuity with most of the chromosomes covered telomere to telomere. Repeat analysis revealed that approximately 61% of the genomes were repetitive sequence. The BUSCO completeness scores ranged from 95.0% to 98.9%, confirming good coverage of the genomes. Gene prediction identified 37198 to 40534 genes. The ks distribution plot of *Macadamia* and *Telopea* suggests *Macadamia* has undergone a whole genome duplication event prior to divergence of the four species and that *Telopea* genome was duplicated more recently. Synteny analysis revealed a high conservation and similarity of the genome structure in all four species. Differences in the content of genes of fatty acid and cyanogenic glycoside biosynthesis were found between the species. An antimicrobial gene with a conserved cysteine motif was found in all four species. The four genomes provide reference genomes for exploring genetic variation across the genus in wild and domesticated germplasm to support plant breeding.

## Introduction

Macadamia, a genus of evergreen trees from the Proteaceae family, is highly valued for its unique flavour, texture, and nutritional properties. It is native to Australia but has now been introduced and widely cultivated in different parts of the world including Hawaii, South Africa, Vietnam, China and Central and South America. *Macadamia* is a genus of four species *M. integrifolia* (Maiden & Betche), *M. tetraphylla* (L. A. S. Johnson), *M. ternifolia* (F. Muell), and *M. jansenii* (C.L. Gross) of which only *M. integrifolia, M. tetraphylla*, and their hybrids are used for commercial production of edible kernel. The other two species are non-commercial due to the high content of cyanogenic glycosides in the mature kernels (Trueman, 2013). Due to the lack of high quality genomic data of *Macadamia,* the crop improvement breeding programs have been based on the phenotypic characteristics, mainly of the two commercial species which risks reducing genetic diversity (O’Connor et al., 2018; Kilian et al., 2021). Several macadamia genomes have been reported recently. The *M. integrifolia* (HAES 741) genome was the first to be sequenced using Illumina short reads (Nock et al., 2016). This 518 Mb assembled genome was highly fragmented (N50 4,745 bp) and incomplete having 77.4% BUSCO genes and covering only 79% of the genome (Nock et al., 2016). HAES 741 was again reassembled using combined Pacific Biosciences (PacBio) long read data along with the Illumina short read sequences (Nock et al., 2020a). A chromosome level assembly was achieved using seven genetic linkage maps This assembly was more contiguous than the previous one with a size of 745 Mb, N50 of 413 kb and 90.2% of BUSCO genes. *M. jansenii* was first de-novo assembled at contig level using three different types of long read sequencing methods (Pacific Biosciences (Sequel I), Oxford Nanopore Technologies (PromethION), and BGI (single-tube Long Fragment Read) for the comparison of the sequencing platforms (Murigneux et al., 2020). All three resulting contig assemblies were highly contiguous and complete, where PacBio continuous long reads (CLR) contig assembly outperformed others in terms of contiguity (N50 1.55 Mb). This PacBio CLR *M. jansenii* contig level assembly was scaffolded to chromosome level using chromosome confirmation capture (Hi-C), where 762 contigs were reduced to 219 scaffolds where 14 scaffolds were of chromosome length, the genome contiguity was improved more than 50 times (N50 52.1 Mb) with 97% BUSCO (Murigneux et al., 2020; Sharma et al., 2021a).

All four *Macadamia* species were sequenced and assembled using the advanced phase assembly (IPA) assembler with PacBio circular consensus sequence (CCS) or HiFi reads for each of the four species. This study reported PacBio HiFi contig level assembly outperformed the earlier CLR contig and scaffold assembly, even with less than half of the volume of sequence data, for *M. jansenii* (Sharma et al., 2021c). A further update on the *M. jansenii* contig level assembly reported the possibility of achieving *de novo* assembly of near chromosome level from sequenced data alone, without using any scaffolding method (Sharma et al., 2022). Recently, a more contiguous and complete assembly of the *M. integrifolia* Chinese cultivar-GUIRE 1(GR1) (Xia et al., 2022) and the *M. tetraphylla* genome were also reported (Niu et al., 2022). The *M. integrifolia* (GR1) chromosome level genome was assembled using Nanopore sequencing, producing a genome of 807 Mb, with a scaffold N50 of 54.7 Mb and 95.7% BUSCO. The *M. tetraphylla* genome was assembled with Hi-C to give a 750 Mb genome, N50 51 Mb, BUSCO of 90%.

The available genome assemblies of macadamia, except *M. ternifolia*, present a challenge for conducting comparative genome analysis due to the use of different sequencing and assembly technologies. To address this limitation, this study aimed to assemble all the genomes of the four *Macadamia* species based upon HiFi sequence data and applying the HiFiasm assembly method. This approach enabling more reliable and accurate comparative genome analysis. The genomic data generated from this study will help in identifying species-specific genes and the variations among the four species. Genes for desirable characteristics present in the non-commercial species may be identified for incorporation into domesticated cultivars, to widen the gene pool of domesticated macadamia.

## Results

### HiFiasm contig assembly

The HiFiasm contig assembly of the four *Macadamia* species resulted in collapsed assemblies that were highly contiguous with N50 more than 45 Mb whereas the haploid assemblies were less contiguous and slightly smaller in size as compared to the collapsed assemblies. The *M. integrifolia* contig assembly had the largest number of contigs, 1049 whereas *M. tetraphylla* had the least. The haploid 1 assembly of all the species was comparatively more contiguous and longer than the haploid 2 assembly (Table S1). The BUCSO analysis revealed a high percentage of genome completeness, with more than 97% coverage. Among the identified BUSCO genes, the majority were found as single-copy genes, with percentages ranging from 83.3% to 84.1%. A small proportion of the BUSCOs were detected as duplicated genes (double BUSCOs), with percentages ranging from 13.4% to 14.2%. Additionally, minor percentage of fragmented BUSCOs in the assemblies, ranging from 0.6% to 0.9% was also reported. The percentage of missing BUSCOs, representing genes absent from the assemblies, was found to be low, varying from 1.4% to 2.6% (Table S1).

### Chromosome level assembly

The Ragtag scaffold assembly length indicated the total size of the genome assemblies for each species, ranged from 735 Mb to 795 Mb. The collapsed assembly was slightly larger than individual haploid assemblies and the Hap2 assembly had the smallest size, ranging from 735 Mb to 776 Mb for each species. Among the species, *M. tetraphylla* had the longest collapsed assembly, while *M. integrifolia* had the shortest. The length of the collapsed assembly for each species reflects the total size of their merged haplotypes, providing a more complete view of their respective genomes. *M. tetraphylla* had the longest haploid assembly, while *M. jansenii* had the shortest. Among the chromosomes in the collapsed genome assemblies of the four species, chr 9 (70 to 75Mb) and chr 10 (68 to 72 Mb) consistently exhibit the greatest lengths. On the other hand, the smallest chromosome in all collapsed assemblies was chromosome 7. The overall BUSCO completeness scores ranged from 95.0% to 98.9%, indicating that a significant proportion of the BUSCOs were present in the assemblies. The majority of BUSCOs were found as single-copy genes, with percentages ranging from 81.6% to 84.2%, confirming the accurate representation of essential genes in the collapsed assemblies. Only a small percentage of BUSCOs appeared as fragmented or missing BUSCO genes, suggesting robust and reliable genome assembly results (Table 1). The N50 values for the collapsed assemblies ranged from 51.7 Mb to 56 Mb. *M. tetraphylla* exhibited the highest N50 values, while *M. ternifolia* had the lowest. These N50 values indicate that the collapsed assemblies have relatively contiguous contigs. The N50 values for the haploid assemblies were generally smaller than those of the collapsed assemblies. The N50 values for the haploid assemblies ranged from 51.4 Mb to 54.8 Mb. The k-mer analysis showed that *M. jansenii* had a smallest genome and low heterozygosity, whereas *M. integrifolia* and *M. tetraphylla* possessed larger genomes and higher heterozygosity. A substantial portion (approximately 63-69%) of their genetic sequences was found to be unique (Table S2.1 & Figure S1a-d). The genome size estimation by flow cytometry results showed *M. tetraphylla* had the largest genome size followed by *M. ternfolia*, which aligns with the assembled scaffolded assembly results (Table S2)

**Table 1:** Chromosome level assemblies of four species of Macadamia representing each chromosome length, BUSCO and N50 values.

|  | *M. jansinii* |  |  | *M. ternifolia* |  |  | *M. integrifolia* |  |  | *M. tetraphylla* |  |  |
| --- | --- | --- | --- | --- | --- | --- | --- | --- | --- | --- | --- | --- |
|  | hap1 | hap2 | collapsed | hap1 | hap2 | collapsed | hap1 | hap2 | collapsed | hap1 | hap2 | collapsed |
| chr_01(Mb) | 54.8 | 56.2 | 57.3 | 55.2 | 58.4 | 58.4 | 56.5 | 58.3 | 58.4 | 59.4 | 57.8 | 59.6 |
| chr_02(Mb) | 49.3 | 44.0 | 51.4 | 50.1 | 48.4 | 50.9 | 45.3 | 46.8 | 47.5 | 48.1 | 49.9 | 50.2 |
| chr_03(Mb) | 50.5 | 50.5 | 52.2 | 50.9 | 51.8 | 51.8 | 50.1 | 50.9 | 51.8 | 51.9 | 50.6 | 51.9 |
| chr_04(Mb) | 56.3 | 51.8 | 56.5 | 61.6 | 55.8 | 62.0 | 56.1 | 57.4 | 58.0 | 57.2 | 63.2 | 63.2 |
| chr_05(Mb) | 47.3 | 47.0 | 48.0 | 47.5 | 46.4 | 47.5 | 47.0 | 45.5 | 47.3 | 45.9 | 46.3 | 47.2 |
| chr_06(Mb) | 54.2 | 53.9 | 54.8 | 53.9 | 51.8 | 53.9 | 52.9 | 53.1 | 53.8 | 54.9 | 56.0 | 56.1 |
| chr_07(Mb) | 45.6 | 42.9 | 46.1 | 46.1 | 44.4 | 46.1 | 46.8 | 44.4 | 44.2 | 44.3 | 43.0 | 44.3 |
| chr_08(Mb) | 47.9 | 48.1 | 48.3 | 48.4 | 47.8 | 48.4 | 46.2 | 50.4 | 50.6 | 51.5 | 52.9 | 53.3 |
| chr_09(Mb) | 70.7 | 70.3 | 71.9 | 76.0 | 70.8 | 70.5 | 72.9 | 73.7 | 75.2 | 73.6 | 74.8 | 76.9 |
| chr_10(Mb) | 71.5 | 65.6 | 71.3 | 62.2 | 64.9 | 72.3 | 63.9 | 64.5 | 68.1 | 71.7 | 63.9 | 71.7 |
| chr_11(Mb) | 57.8 | 57.6 | 59.3 | 60.8 | 61.4 | 63.9 | 61.5 | 60.7 | 61.9 | 61.3 | 64.6 | 63.5 |
| chr_12(Mb) | 49.2 | 48.3 | 49.2 | 50.2 | 44.5 | 50.2 | 47.7 | 41.0 | 47.8 | 49.6 | 47.4 | 49.6 |
| chr_13(Mb) | 49.9 | 46.0 | 49.9 | 47.3 | 49.2 | 49.1 | 47.5 | 47.3 | 49.7 | 48.2 | 47.6 | 48.7 |
| chr_14(Mb) | 56.7 | 53.2 | 57.1 | 55.9 | 53.0 | 55.9 | 53.6 | 57.8 | 61.2 | 58.8 | 57.7 | 58.9 |
| Assembly Length | 761 Mb | 735 Mb | 773 Mb | 766 Mb | 748 Mb | 780 Mb | 748 Mb | 751 Mb | 775 Mb | 776 Mb | 775 Mb | 795 Mb |
| Complete BUSCO | 98.9% | 95.0% | 97.7% | 97.1% | 96.5% | 97.7% | 95.1% | 94.3% | 97.6% | 97.4% | 97.3% | 97.8% |
| Single | 83.3% | 82.1% | 84.2% | 83.8% | 83.4% | 84.1% | 82.4% | 81.6% | 84.1% | 83.5% | 83.8% | 83.7% |
| Double | 13.6% | 12.9% | 13.5% | 13.3% | 13.1% | 13.6% | 12.7% | 12.7% | 13.5% | 13.9% | 13.5% | 14.1% |
| Fragmented | 0.6% | 0.6% | 0.7% | 0.8% | 0.8% | 0.8% | 0.9% | 0.6% | 0.6% | 0.8% | 0.7% | 0.7% |
| Missing | 2.5% | 4.4% | 1.6% | 2.1% | 2.7% | 1.5% | 4.0% | 5.1% | 1.8% | 1.8% | 2.0% | 1.5% |
| N50 | 54.2 Mb | 51.7 Mb | 54.7 Mb | 53.8 Mb | 51.8 Mb | 53.8 Mb | 52.8 Mb | 53 Mb | 53.7Mb | 54 Mb | 56 Mb | 56 Mb |

\*The chromosomes were numbered according to the *M. integrifolia* genome (Nock et al., 2020b) which used the seven genetic linkage maps.

### Genome structure comparison

The genomic structure comparison of the four *Macadamia* species using SyRI revealed syntenic regions, inversions, translocations, and duplications. Chromosomes 9 and 10 showed several structural rearrangements, with chr 9 exhibiting changes in the first half and chr 10 in the second half. Chr 04 also displayed genomic rearrangements at one end, while chr 12 in all four species showed several duplications in the middle (Figure 1). Dotplots of the reference genome (*M. jansenii* Hi-C) against the four *Macadamia* species (assembled by ragtag) showed varying structural rearrangements, with *M. integrifolia* and *M. tetraphylla* having more structural differences compared to *M. jansenii* (Figure S3). Among all chromosomes, chr 9 and 10 had the majority of rearrangements. Similarly, dotplot comparison between the haploid assemblies showed *M. integrifolia* haploids were the most diverse, while *M. jansenii* haploids were the least diverse (Figure S3). The study showed that the genomes of different *Macadamia* species have different structures and arrangements, showing their unique genetic characteristics.

**Figure 1:**
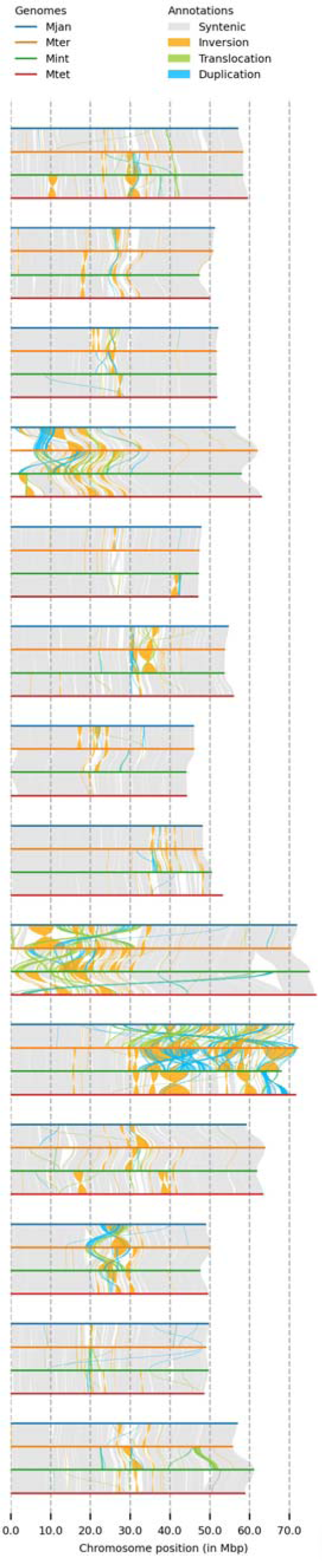
The genome structure comparison of four *Macadamia* species, with different colours denoting each species and structural rearrangements (synteny, inversion, translocation, and duplication) as indicated on the top of the image.

### Genome annotation

The repeat content analysis of the four species identified total 61% to 62% across both haploid and collapsed assemblies. This indicates that a major portion of the genomes is composed of repetitive elements. Among the different repeat types, Long Terminal Repeat (LTR) elements were the most prevalent, comprising around 22.1% to 23.8% of the genomes, followed by Long interspersed nuclear elements (LINE) elements. Other repeat types, such as DNA elements, unclassified elements, small RNA elements, satellites, and simple repeats, contributed to a smaller fraction of the total repeat content, ranging from 4.13% to 6.51% (Table S3). The consistency of the total repeat content between haploid and collapsed assemblies suggests that the repetitive landscape is preserved even after haplotype merging. Comparing the collapsed assemblies with their respective haplotypes, for the number of predicted genes, it was observed that the gene content remained relatively stable. Among the collapsed assemblies, *M. integrifolia* exhibit the highest number of genes, 40534 while *M. jansenii*, exhibit lowest number of genes, 37198. In the haploid assemblies, the number of genes ranges from 36465 to 47388. The number of genes distribution across the chromosomes, showed chr 09 and 10 have more genes than the other chromosomes (Table 2). The higher number of CDS and protein sequences identified by Braker3 compared to the gene count is because some genes produce multiple transcripts through alternative splicing. The telomere analysis revealed that the collapsed assemblies generally exhibited “telomere to telomere” arrangements for most chromosomes. However, a few exceptions were observed, where telomere was present only at one of the ends, suggesting missing or ambiguous telomeric sequences on some chromosome ends (Table S4). The functional annotation of the CDS sequences, showed majority of the similarity hits with *Telopea*, the only other member of the Proteaceae with a high-quality genome sequence. All the species showed similarity with *Telopea* followed by *Nelumbo nucifera* and *Tetracentron sinense* (Figure S4). The pathway analysis of the annotated CDS sequences, identified a consistent number of pathways among the four species, *M. jansenii* and *M. tetraphylla* each identified 580 pathways, 578 pathways in *M. ternifolia* and *M. integrifolia* exhibited 581 pathways. The top five pathways, namely purine and thiamine metabolism, response to drought, biosynthesis of cofactors, and starch and sucrose metabolism, were found in all four species. This suggests that these pathways play crucial roles in the biological processes and responses shared by all four species.

**Table 2:** Distribution of genes across the 14 chromosomes of Macadamia species.

|  | *M. jansanii* |  |  | *M. ternifolia* |  |  | *M. integrifolia* |  |  | *M. tetraphylla* |  |  |
| --- | --- | --- | --- | --- | --- | --- | --- | --- | --- | --- | --- | --- |
|  | Hap 1 | Hap2 | Collapsed | Hap 1 | Hap2 | Collapsed | Hap 1 | Hap2 | Collapsed | Hap 1 | Hap2 | Collapsed |
| Chr_01 | 2483 | 2543 | 2474 | 2455 | 2484 | 2612 | 2483 | 2389 | 2665 | 2643 | 2521 | 2631 |
| Chr_02 | 2666 | 2514 | 2608 | 2774 | 2666 | 2739 | 2453 | 2613 | 2699 | 2664 | 2735 | 2786 |
| Chr_03 | 2802 | 2868 | 2844 | 3007 | 2943 | 3053 | 2837 | 2771 | 2974 | 2949 | 2917 | 3017 |
| Chr_04 | 2780 | 2670 | 2718 | 2832 | 2706 | 2931 | 2833 | 2746 | 3078 | 3142 | 2782 | 2813 |
| Chr_05 | 2800 | 2783 | 2798 | 2798 | 2636 | 2911 | 2755 | 2569 | 2814 | 2746 | 2780 | 2866 |
| Chr_06 | 2607 | 2579 | 2568 | 2623 | 2465 | 2683 | 2585 | 2616 | 2667 | 2702 | 2731 | 2709 |
| Chr_07 | 2790 | 2702 | 2696 | 2764 | 2699 | 2836 | 2810 | 2587 | 2623 | 2711 | 2578 | 2712 |
| Chr_08 | 2768 | 2768 | 2677 | 2742 | 2671 | 2770 | 2509 | 2802 | 2878 | 3062 | 2869 | 2837 |
| Chr_09 | 2870 | 2897 | 2878 | 2915 | 2874 | 3053 | 3373 | 2816 | 3842 | 3626 | 2978 | 3137 |
| Chr_10 | 2402 | 2359 | 2428 | 2301 | 2209 | 2463 | 2699 | 2367 | 3103 | 3710 | 2295 | 2392 |
| Chr_11 | 2820 | 2896 | 2812 | 2910 | 2845 | 3001 | 2917 | 2879 | 3087 | 2888 | 3024 | 2935 |
| Chr_12 | 2590 | 2567 | 2517 | 2642 | 2408 | 2721 | 2576 | 2092 | 2538 | 2617 | 2430 | 2566 |
| Chr_13 | 2766 | 2627 | 2732 | 2641 | 2716 | 2790 | 2684 | 2723 | 2875 | 2694 | 2663 | 2724 |
| Chr_14 | 2560 | 2409 | 2448 | 2598 | 2474 | 2626 | 2446 | 2495 | 2691 | 2634 | 2534 | 2608 |
| Total no. of genes | 37704 | 37182 | 37198 | 38002 | 36796 | 39189 | 37960 | 36465 | 40534 | 40788 | 37837 | 38733 |
| Number of mRNA | 43510 | 43098 | 43092 | 44506 | 43016 | 45694 | 44527 | 43010 | 47301 | 47184 | 44490 | 45519 |
| Number of CDS | 43510 | 43098 | 43092 | 44506 | 43016 | 45694 | 44527 | 43010 | 47301 | 47184 | 44490 | 45519 |

### Gene family analysis

#### Anti-microbial gene analysis

The homologs of an anti-microbial gene was identified in all four species of *Macadamia* by using a BLAST search. Only one gene was identified in all four species on chr 9. The sequence alignment of the reference gene MiAMP-2 with copies in all four species, revealed a high degree of homology (Figure S5). This protein sequence alignment clearly shows four repeated segments with four a cysteine motif C-X-X-X-C-(10±12)-X-C-X-X-X-C.

### Fatty acid pathways

The number of FatA and FatB genes, essential for fatty acid production, varied between species. *M. integrifolia* had the highest number of both genes, 10 and 11, respectively, suggesting the potential of this species for robust fatty acid synthesis. SAD (Stearoyl-ACP Desaturase) genes, which are mainly responsible for converting stearic acid (C18:0, SA) to oleic acid (C18:1, OA) (Si et al., 2023), were present in high numbers across the four species, indicating their active involvement in the desaturation processes. This supports the observations of Hu et al., (2022). The conversion of C16:0 to C18:0 through elongation is a more efficient process compared to the conversion of C16:0 to C16:1 and the desaturation of C18:0 to C18:1 appears to be more effective than the desaturation of C16:0 to C16:1 (Hu et al. (2022). KAS (Ketoacyl-ACP Synthase) genes, crucial for fatty acid chain elongation, are notably absent in *M. integrifolia*, potentially indicating a unique fatty acid metabolism pathway in this species. In contrast, the other three species possess KAS genes, particularly *M. jansenii* and *M. ternifolia* (10 each), highlighting their capacity for elongating fatty acid chains (Table S5 (A)).

### Cyanogenic glycoside pathway

CYP 79 which catalyse the first step in the biosynthesis of cyanogenic glycosides by acting on amino acids and converting them into aldoximes (Irmisch et al., 2013) was found to be present in *M. integrifolia* and *M. tetraphylla* and absent in *M. jansenii* and *M. ternifolia*, indicating a potential deviation from the typical cyanogenic glycoside biosynthesis pathway in these species. In contrast, CYP71, responsible for further converting aldoximes into cyanohydrin (Hansen et al., 2018), was uniformly present among all the species. The number of BGLU and UGT genes, which are responsible for the detoxification and the glycoside modification was found to vary across the four species, reflecting differences in detoxification capabilities in the cyanogenic pathway. *M. tetraphylla* lacks UGT genes entirely, potentially indicating unique detoxification mechanisms (Table S5 (B)).

### WRKY genes

The WRKY gene family, known for its key role in plant development and stress responses (He et al., 2019), revealed varying protein counts ranging from 58 to 61 among the four *Macadamia* species (Table S5 (C)). These findings align with the prior discovery of 55 WRKY proteins within the *M. tetraphylla* genome as reported by Niu et al. in 2022.

### Orthologous and Phylogenetic analysis

Orthologous clusters were generated across the four *Macadamia* species using *Telopea* as the outgroup, to identify genes that have been conserved across different species and may have similar functions. The clustering patterns of gene families across five plant species: *T. speciosissima* and the four *Macadami*a species revealed a total of 195004 proteins grouped into 34696 gene clusters. Among all the clusters only 31 clusters showed overlaps among two or more of the plant species and 8217 single-copy clusters indicated conserved genes among the five species (Table S6). A total of 30111 (15.4%) singleton or species-specific gene were found in 2090 unique gene clusters, where *Telopea* contains the maximum number of unique gene clusters (902). Among the *Macadamia* species, *M. integrifolia* had the maximum (403) whereas *M. jansenii* the lowest number of singleton gene clusters (201) (Figure 2 & Figure S3). The Gene Ontology (GO) enrichment analysis of these unique gene clusters holds great promise in providing valuable insights into the distinct biological functions and potential adaptations of each species.

**Figure 2:**
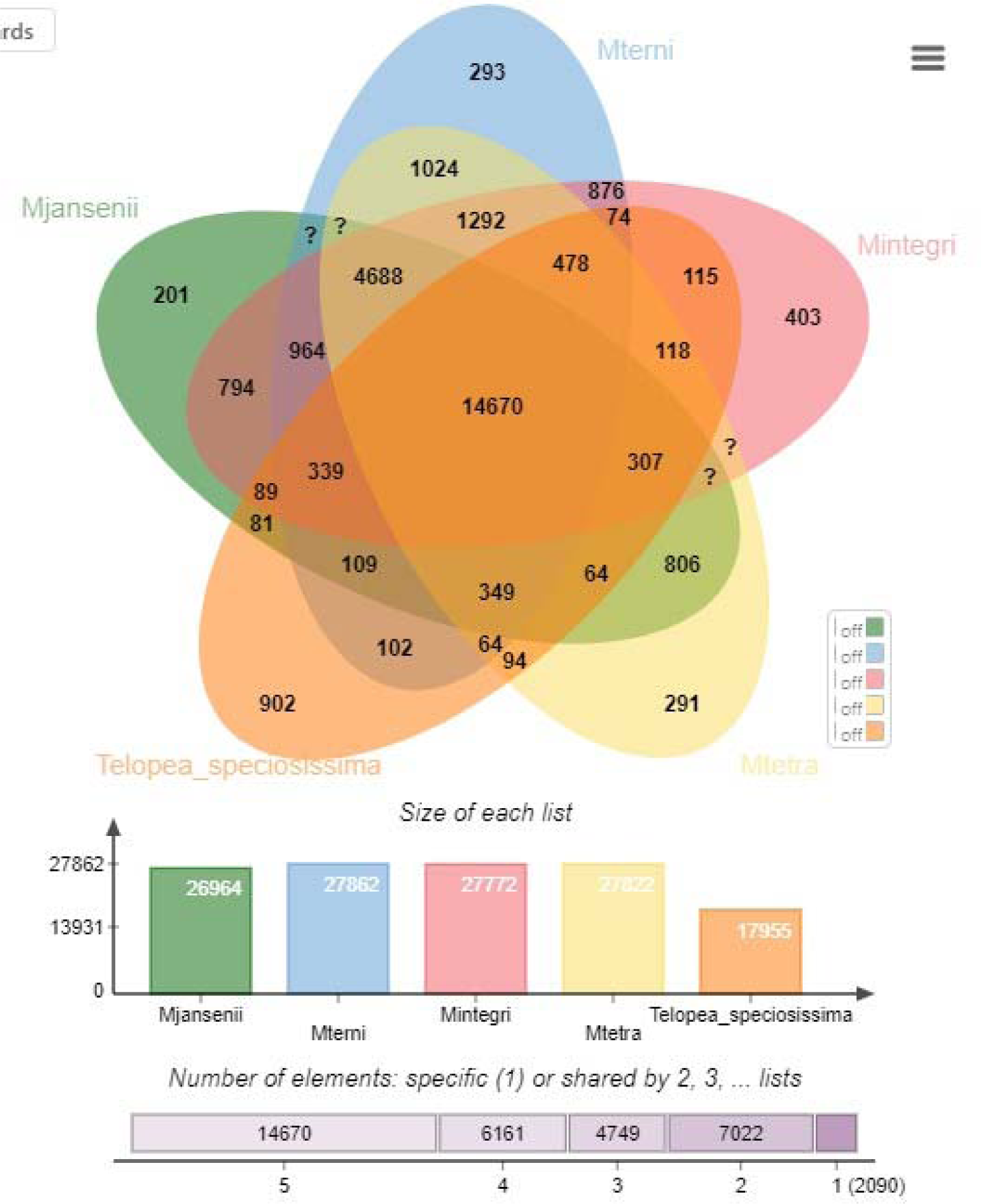
A Venn-diagram showing clusters of orthologous groups of genes (OGs) for the four *Macadamia* species and *T. speciosissima*. Number of orthologous groups (OGs) belonging to core genome (OGs common among all five species-union of all circles), number of singletons (unique genes—outer area of each circle), and the common ones of remaining different combination of all five species (in between the core and the periphery of the diagram) are described.

A phylogenetic tree was constructed to investigate the genetic divergence and evolutionary distances among the *Macadamia* species, with *Telopea* as the outgroup. The tree indicates two main branches. One branch includes *M. integrifolia* and *M. tetraphylla*, indicating a shared genetic lineage. The other branch comprises *M. jansenii* and *M. ternifolia*, highlighting their distinct genetic lineage. (Figure S6).

### WGD and Synteny

The analysis of ks values in all four species of *Macadamia* genomes revealed a distinctive peak at ks≈ 0.32 (Figure 3). The *Telopea* genome exhibited a peak at ks≈ 0.28. This comparison of the peaks in *Macadamia* and *Telopea* suggests a more recent whole-genome duplication (WGD) event in *Telopea* compared to *Macadamia*. In some WGD studies, WGD and divergence time estimation have been based solely on ks values. However, in recent years, there has been growing research cautioning against exclusively relying on ks plot analysis for these estimations. Instead, additional sources of evidence are recommended to ensure a more robust WGD assessments (Tiley et al., 2018, Zwaenepoel et al., 2019).

**Figure 3:**
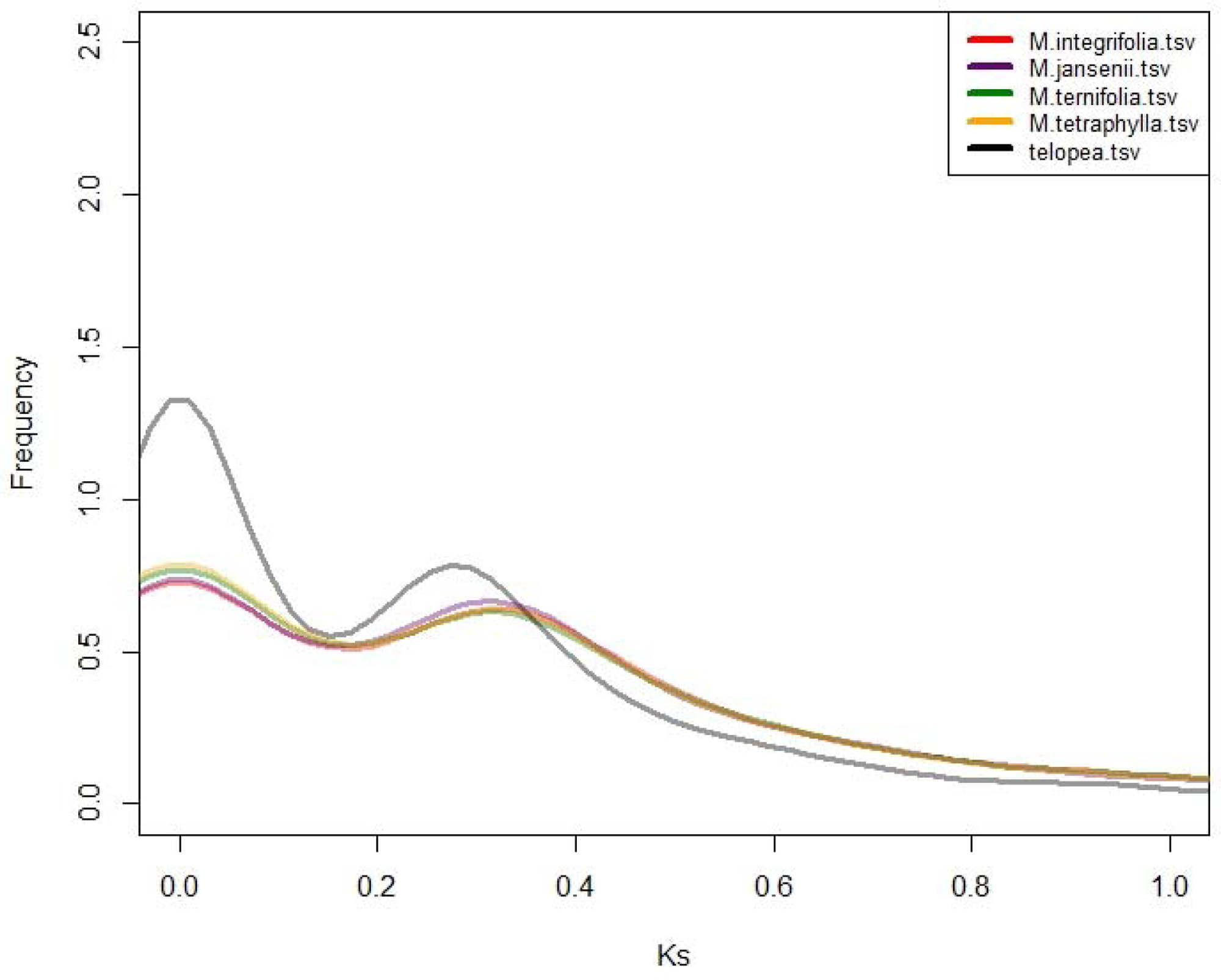
Ks distribution plot of the four *Macadamia* species and *Telopea*. The colour code of each species is provided on the top left corner.

The duplication events were further verified using the synteny plots which highlighted the duplicated genetic regions and genes. Synteny analysis revealed extensive genetic similarity within the species and among the four species, particularly on chromosomes 9 and 10 (Figure 4 & Figure S8)

**Figure 4:**
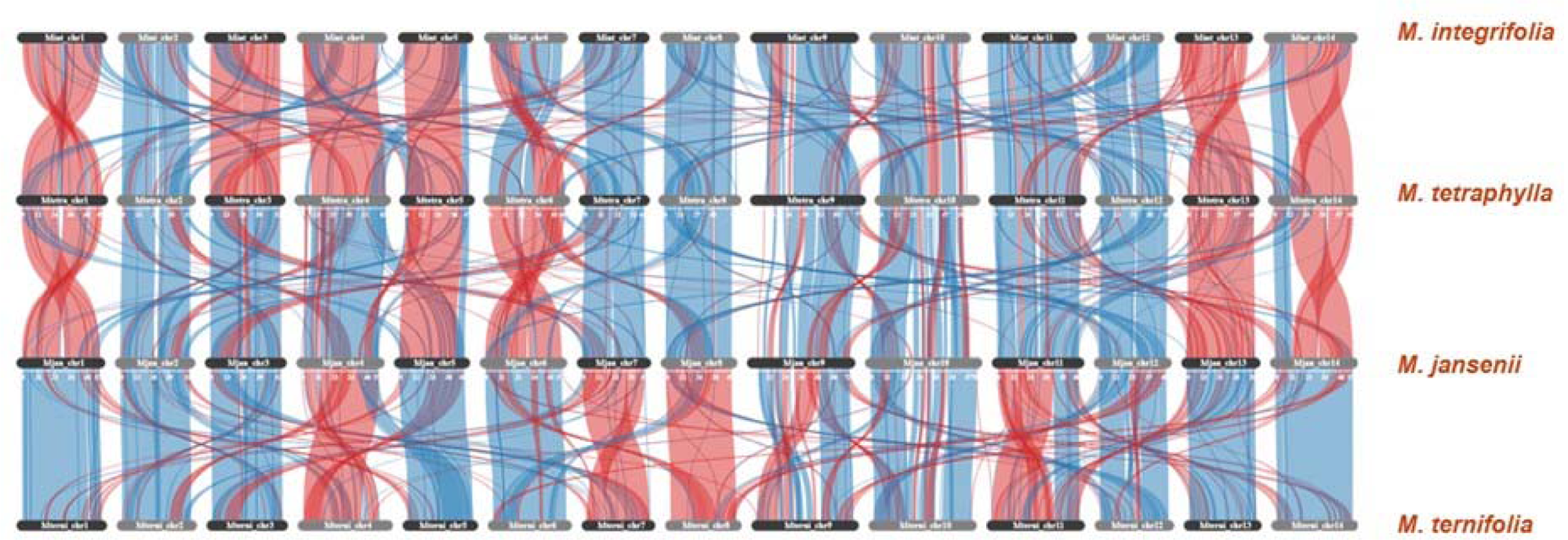
Synteny plot across all the four *Macadamia* species. The vertical lines connect orthologous genes across the four species. The blue coloured ribbons represent the regular conserved regions while the red ribbons represent the inverted regions.

### Expansion-contraction of gene families

The study of differences in protein families among the annotated species revealed significant differences between the groups. The protein family size varied notably between the *Macadamia* species and *Telopea*. A total of 613 different protein clusters were contracted and only 21 protein family clusters showed expansion in *Macadamia* as compared to *Telopea*. Among the two clades of *Macadamia*, the edible, species (*M. integrifolia* and *M. tetraphylla*) exhibited more expansion-contraction (+18/-140) than the bitter non-edible species (+0/-5) (Figure 5). Among 5 contracted clusters of the bitter species, one cluster belonged to Xanthotoxin 5-hydroxylase CYP82C4, which is expressed in roots under iron-deficient conditions.

**Figure 5:**
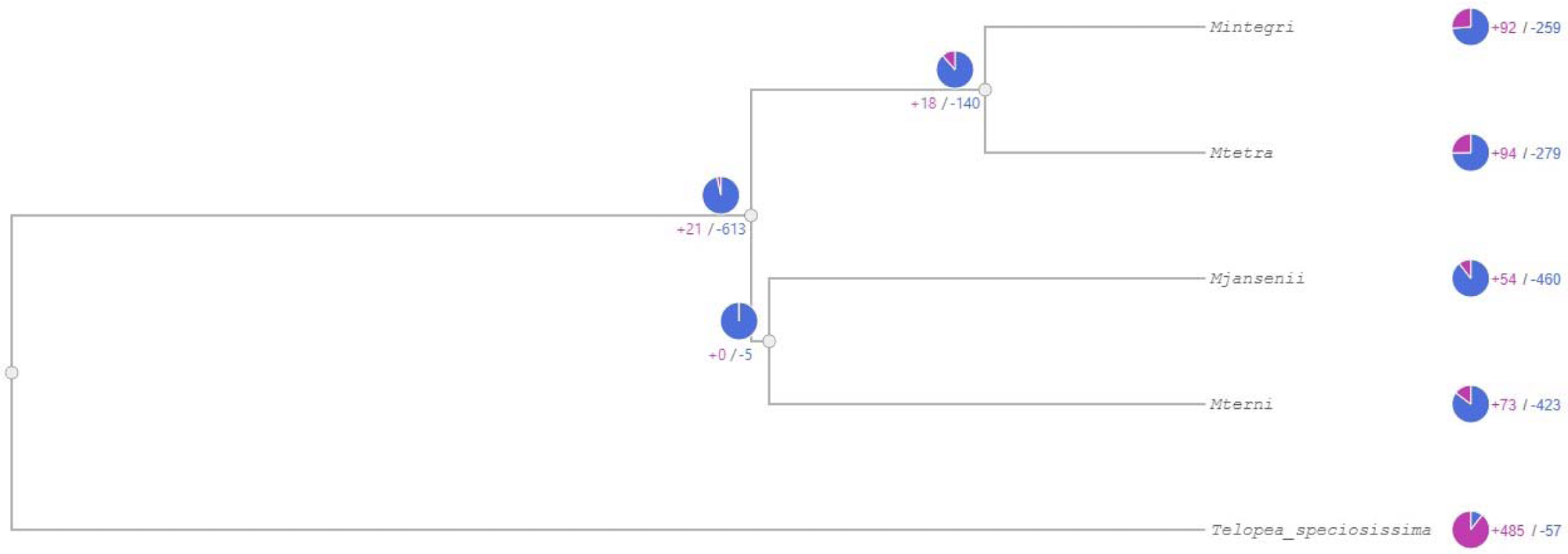
Gene family Expansion and contraction across the *Macadamia* species and *Telopea*. The blue colour represents contraction and pink presents expansion of gene clusters.

All the four species of *Macadamia* individually displayed more contraction than expansion. The expansion ranging from 259 to 423 clusters of protein, where *M. jansenii* showed the highest number of contractions, followed by *M. ternifolia,* and *M. tetraphylla*. Whereas only 54-94 protein clusters were expanded, and M. tetraphylla displayed the highest expansion of proteins (+94), one of these expanded clusters was associated with the GO term ‘rejection of self-pollen’ However, for *Telopea* the opposite was found with more expansion than contraction (+485/-57) of protein clusters (Figure 5). Both the edible species shows similar changes and the gene enrichment analysis of both also showed similar pattern, and the same held true for the non-edible species.

## Discussion

In this study, a high-quality reference genome and annotations were created for the four species of *Macadamia*. The gene model set completeness, as measured by BUSCO, suggested that the annotation pipeline used was suitable for comprehensive capture of protein-coding genes. The comparison of genome assemblies of the already available genomes of *M. jansenii*, *M. integrifolia,* and *M. tetraphylla* with those generated in this study revealed notable improvements in the assembly statistics. For *M. jansenii*, the newly assembled genome demonstrated an increase in length (from 758Mb to 773Mb), improvement in N50 value from 52Mb to 55Mb and slight improvement in BUSCO as compared to the already available *M. jansenii* Hi-C assembly’s 758 Mb (Sharma et al., 2021b). This study has greatly improved the *M. integrifolia* (cultivar 741) genome with a longer assembly length of 775 Mb and a significantly higher BUSCO of 97% and N50 value of 53 Mb as compared to previous assemblies by Nock et al., in 2016 (N50: 4.7 kb) & 2020 (N50: 413 kb) (Nock et al., 2016; Nock et al., 2020b). Similarly, the *M. tetraphylla* genome showed great improvement in terms of N50 56 Mb and 98% BUSCO as compared to already available *M. tetraphylla* genome (Niu et al., 2022). The genome assemblies generated in this study provide enhanced continuity, higher BUSCO completeness, and increased gene identification compared to previous versions, providing a robust basis for genome comparison. Additionally, the genome assemblies attained complete chromosome coverage from telomere to telomere for most of the chromosomes, which has not been reported in the previous studies.

The comparison of collapsed assembly statistics of four *Macadamia* species revealed *M. tetraphylla* assembly stood out with the longest genome length. The *M. jansenii* has the shortest assembly length among the four. The gene content comparison across the four species revealed that *M. integrifolia* assembly exhibited the highest number of genes, followed by *M. ternifolia* and *M. tetraphylla*. These variations in gene counts may be attributed species-specific genomic features. Haploid-resolved assemblies are essential in genomics research, as they facilitate accurate gene phasing, improved annotation, and enhanced insights into genetic diversity (Nakandala et al., 2023; Zhang et al., 2021; Cheng et al., 2021). Heterozygosity between the haplotypes in diploids can complicate the genome assemblies. The low heterozygosity of *M. jasnenii* and high heterozygosity of *M. integrifolia* and *M. tetraphylla* (Sharma et al., 2021b; Xia et al., 2022; Nock et al., 2020b; Niu et al., 2022) was also supported by k-mer analysis, haploid assembly statistics and dotplot comparisons. The dotplot comparison of the two *M. jansenii* haploid assemblies, showing minimal differences between the two. On the other hand, the highly heterozygous species, *M. integrifolia* and *M. tetraphylla*, exhibit significant differences in the dotplots, gene numbers, structural rearrangements and individual chromosome lengths. These findings highlight the genomic variations at haploid levels among the different *Macadamia* species, providing valuable insights into their genetic diversity.

Antimicrobial proteins (AMP) are essential components of plant innate immunity, exhibiting diverse activities such as antibacterial, antifungal, insecticidal, and antiviral effects, enabling effective defense against pathogens and pests (McManus et al., 1999; Li et al., 2021a). Comparative analysis of AMP protein across the four macadamia species, showed that the gene location remained conserved on chr 9 across all the species and the sequence alignment revealed a highly conserved eight motif pairs of cysteines, however the amino acid sequence was variable. These results aligned with (Li et al., 2021b; McManus et al., 1999; Campos et al., 2018). The variable distribution of CYP79, across the four species, may indicate potential deviations from the conventional cyanogenic glycoside biosynthesis pathway in the two bitter species, *M. jansenii* and *M. ternifolia.* In contrast, CYP71’s uniform distribution across all species, indicating its essential role. The differential counts of detoxifying enzymes, BGLU and UGT, underscore species-specific strategies, with lack of UGT genes in *M. tetraphylla* suggesting a different detoxification mechanism. The analysis of fatty acid pathway genes showed *M. integrifolia* stands out prominently with the highest counts for both FatA and FatB genes, signifying its robust capability for fatty acid production and may explain the domestication of *Macadamia* being based mainly on this species. Additionally, the higher abundance of SAD genes across the four species suggests their active role in desaturation, as confirmed by Hu et al. (2022), highlighting the efficiency of C18:0 to C18:1 conversion. The absence of KAS genes in *M. integrifolia* suggests a potential uniqueness in its fatty acid metabolism pathway, distinct from the other three species, which possess KAS genes (especially *M. jansenii* and *M. ternifolia* with 10 each), highlighting their capacity for extending fatty acid chains. Variations in WRKY protein counts (ranging from 58 to 61) across *Macadamia* species supporting their roles in development and stress responses.

Utilizing long-read assemblies in this study of *Macadamia* gene families significantly increased the accuracy of results for expansion and contraction events. This accuracy is crucial for identifying essential genes and gene families involved in important biological processes and hence the accurate interpretation of expansion-contraction (CAFE) analysis. Remarkably, the edible macadamia species demonstrated a higher incidence of expansion-contraction, while the bitter species exhibited fewer changes. This observation implies potential differences in the distribution of gene families between the two groups, suggesting distinct evolutionary trajectories. Understanding the factors behind the expansion of particular gene families in edible *Macadamia* species could provide valuable clues about the evolution of *Macadamia* and be harnessed for the development of improved cultivars with desirable traits. Moreover, the presence of common ks peaks events in the four *Macadamia* species suggests significant evolutionary events that have shaped their genomes. Comparison of the ks plot between the *Macadamia* and the *Telopea* genomes, suggests that *Telopea* has undergone a more recent duplication event as compared to *Macadamia*, though the exact dates of divergence and duplication will require more analysis. Synteny analysis further highlights the conservation of genetic regions and genes within each species and reveals intriguing similarities among the different species, particularly on chromosomes 9 and 10. These findings emphasize the importance of whole genome duplication in shaping the genetic landscape of macadamia and provide valuable insights into the evolutionary dynamics of this economically important crop. The analysis of orthologous clusters and gene families among the four *Macadamia* species and *Telopea* provided valuable insights into the conservation and divergence of genes in these plants. Among the 195,004 proteins grouped into 34,696 gene clusters, only 31 clusters showed overlaps among two or more species, while 8,217 clusters contained conserved single-copy genes across the five species. These unique gene clusters hold great promise for uncovering distinct biological functions and potential adaptations of each species. The phylogenetic tree, with *Telopea* as the outgroup, demonstrates two main branches: one containing *M. integrifolia* and *M. tetraphylla* and the other comprising *M. jansenii* and *M. ternifolia*, illustrating the genetic relationships among the *Macadamia* species. The core orthologous genes, as expected included gene families related to categories like cell growth, DNA replication and repair, metabolism, and cell cycle regulation.

The comparative genomics and experimental study, presented here, allows for the first time a genus-wide view of the biological diversity of the *Macadamia*, which provides a strong foundation for the genome wide analysis.

## Material and Methods

### DNA and RNA sample

The HiFi sequencing data of the four *Macadamia* species (Sharma et al., 2021b) was used for this study. RNA sequence data for *M. jansenii* was used from Sharma et al., 2021a. Total RNA *M. ternifolia* and *M. tetraphylla* was extracted from fresh leaf tissues using Rubio-Pina et al RNA isolation method (Rubio-Piña and Zapata-Pérez, 2011) along with Qiagen kit method and sent for short read sequencing at Macrogen Oceania. RNA Seq data for young leaves of *M. integrifolia* (HAES 741) was downloaded from NCBI SRA data SRR10897159.

### Genome assembly

The HiFi reads of four species were assembled using HiFiasm to generate both the collapsed and the haploid assemblies (Cheng et al., 2021; Sharma et al., 2021c). The contig assembly generated from HiFiasm was then scaffolded using a reference-guided approach with the RagTag tool (Alonge et al., 2019) using *M. jansenii* Hi-C as the reference (Sharma et al., 2021a). The chromosomes were numbered according to the *M. integrifolia* genome (Nock et al., 2014). The contigs more than 1 Mb in size were used as input for the reference guided approach. To assess the completeness of the assembles, the Benchmarking Universal Single-Copy Orthologs (BUSCO) (version v5.4.6) (Simao et al., 2015) was used with the eudicots_odb10 dataset. The genome completeness was evaluated using the quality assessment tool QUAST (Gurevich et al., 2013).

### Genome estimation (flowcytometry and k-mer) and dotplots

For flow cytometry methods nuclei were extracted from leaf tissue by mechanical dissociation as described by Galbraith *et al*. (Galbraith et al., 1983) with modifications for woody plant species. Briefly, 40 mg of young macadamia leaf were co-chopped with 15 mg of the internal standard *Oryza sativa* ssp. Japonica cv. Nipponbare, in 0.4mL of ice-cold nuclear isolation buffer in a 5cm polystyrene Petri dish. For *M. tetraphylla* and *M. integrifolia*, Arumuganathan and Earle (Arumuganathan and Earle, 1991) nuclear isolation buffer was used; while MB01 (Sadhu et al., 2016) nuclear isolation buffer was used for *M. ternifolia* and *M. jansenii*. Samples were chopped for approximately 10-12 minutes, first into fine longitudinal strips with new parts of a sharp razor blade and then into perpendicular slices. Resulting homogenates were gently filtered through a pre-soaked 40-µm nylon mesh into a 5mL round bottom polystyrene tube. Homogenates were then stained with 50µg/mL of propidium iodide (PI) (Sigma, P4864-10ML) and 50µg/ml of RNase A (Qiagen, 19101) for 10 minutes on ice. The BD Biosciences LSR II Flow Cytometer and FlowJo software package was used to analyse the homogenates. Briefly, fluorescence was collected using a 488nm excitation laser tuned to 514.4nm and a 610/20nm bandpass filter. Instrument settings were kept constant across and throughout experiments: forward scatter voltage at 199, side scatter voltage at 300, fluorescence intensity voltage at 500, with a slow flow rate (20-50 events/s). Three biological replicates were performed on three different days. For each biological replicate, a minimum of 1,500 PI-stained events were collected per PI-stained peak. Nuclear DNA content was calculated as previously described (Doležel et al., 2007) using 388.8 Mb at 1C for the assumed size of *O. sativa (Sasaki and International Rice Genome Sequencing, 2005)*.

Genome estimation using K-mer analysis was performed by Jellyfish’s Version 2.3.0 (Marçais and Kingsford, 2011) count and histo commands. The histo file was visualised in genomescope (Ranallo-Benavidez et al., 2020). Dotplots for the assembly comparisons were plotted using the Chromeister (Pérez-Wohlfeil et al., 2019) tool available at Galaxy Australia (https://usegalaxy.org.au/).

### Genome annotation

The identification and classification of the *de novo* repeat elements in all the collapsed assemblies of all four species was performed using the RepeatModeler (version 2.0.2a) (http://www.repeatmasker.org/RepeatModeler/). The repeats identified were then masked by repeatmasker (version 4.0.9) (http://www.repeatmasker.org/). The gene models in the masked assemblies were identified using an *ab-initio* method along with RNA-seq evidence Braker3 version 3.0.3 (Brůna et al., 2021). To prepare the input files for the Braker3 run, the masked assemblies were first aligned with RNA-seq using HISAT2 version 2.1 (Kim et al., 2019), then the output aligned .sam file was converted to a .bam file using samtools (Li et al., 2009). The softmasked genome assembly file along with the sorted bam file was used as input files for the Braker3 pipeline. The protein and coding sequence (CDS) fasta files generated from Braker3 contain multiple transcripts therefore a python script was used to keep only one transcript per gene. The filtered protein and CDS fasta was then used for the downstream analysis. Tidk version 0.2.31 (Telomere identification toolkit) tool (https://github.com/tolkit/telomeric-identifier) was used to identify the telomere region in the genome assemblies using ‘search’ and ‘plot’ commands.

Functional annotation of the gene set identified for each of the four genomes was performed through Omicx box (version 3.0.27) (OmicsBox, 2019). This pipeline consists of BLAST2GO (Conesa and Götz, 2008) and Interproscan (Jones et al., 2014). For BLAST2GO, the ‘blastx-fast’ feature was used with NCBI non-redundant protein sequences (nr v5) database and the e-value was set at 1e-10 with 10 blast hits. The taxonomy filter was set at 33090 Viridiplantae. For Interproscan all the available databases such as families, structural domains, sites and repeats databases were selected. For the pathway analysis: Plant reactome (Gramene) (Naithani et al., 2020) and KEGG pathway (Kanehisa and Goto, 2000) was performed using Omics box.

Gene family analysis: Anti-microbial genes were identified across the four species by conducting a BLAST homology search, looking for transcripts resembling *M. integrifolia’s* antimicrobial cDNA (MiAMP2). Sequence alignment using Clone Manager ver 9.0 was performed with alignment parameter scoring matrix of Mismatch (2), Open Gap (4), and Extension-Gap (1). To identify genes involved in cyanogenic glycoside, fatty acid metabolism and WRKY gene across the four genomes, BLAST was performed and the top hits based on sequence similarity was selected.

### Orthologous and Phylogenetic analysis

Orthologous and phylogenetic analysis was performed using Orthofinder (V2.5.5) (Emms and Kelly, 2019) using the protein sequences of all the four *Macadamia* species along with data for Telopea. The common and unique set of orthologous protein sequences among the five species were plotted using the UpSet plot and the venn diagram of the Orthovenn3 (Sun et al., 2023). The core or single copy orthologs obtained from Orthofinder were used to construct the phylogenetic tree using Orthovenn3.

### Whole genome duplication

Whole genome duplication (WGD) analysis was performed to compute the whole set of paralogous genes in the genome using WGD tool version 1.1.2 (Zwaenepoel and Van de Peer, 2019). Ancient WGDs was calculated by examining the distribution of synonymous substitution per site (Ks) within a genome or Ks distribution. WGD analysis of all the four species of *Macadamia* was performed to estimate the origin and diversification. Wgd ‘dmd’ and ‘ksd’ commands were used to generate the Ks distribution plot.

### Conservation of gene order and genomic regions

A pairwise whole-genome comparison was performed using SyRI (Goel et al., 2019) to find the structural and sequence differences between the two genomes. The genomes were first aligned using the minimap2 (Li, 2018) and samtool (Li et al., 2009) was used to index the alignment BAM file. The BAM file was then used to run the SyRI tool, the same output file was then passed through the visualisation tool plotSR (Goel and Schneeberger, 2022) using default parameters to visualise the synteny and the structural rearrangements between the *Macadamia* species.

### Collinearity and Expansion-contraction of gene families

The degree of collinearity within and between the genomes of the four *Macadamia* species were calculated by using MCScanX (Wang et al., 2012). The protein fasta file of all the four species were combined together and used as input for the all-vs-all homology search with the Blastp algorithm with e-value set at 1e-10, max target sequences at 5 and output format 6. The resulting tabular blastp file along with combined gff file was then fed into MCScanX using default parameters. For self synteny MCScanX was run with default settings with the blastp output and the gff file of individual species. The web based tool - SynVisio (Bandi and Gutwin, 2020) was used to visualize collinearity. The CAFE5 tool of Orthovern3 was used to perform the expansion and contraction of the gene families. All default parameters were used.

## Data availability

The genome sequencing data from PacBio has be submitted under NCBI bioproject PRJNA694456. The genome assemblies and annotation of four *Macadamia* species have been deposited under in Genome warehouse under the bioproject: PRJCA020274.

## Acknowledgements

This project was funded by the Hort Frontiers Advanced Production Systems Fund as part of the Hort Frontiers strategic partnership initiative developed by Hort Innovation, with co-investment from The University of Queensland, and contributions from the Australian Government. We thank the Research Computing Centre (RCC), University of Queensland for support and providing high performance computing resources. We are also thankful to Virginia Nink and the Queensland Brain Institute Flow Cytometry Facility for technical assistance with flow cytometry.

## Contributions

Contributions of authors were as follows: Designed and supervised the project: RJH, AKM, AF, BT and CN. Genome assembly, annotation and downstream analysis: PS and AKM. Flow-cytometry analysis: LC. RNA data: CN. Drafted the manuscript: PS and LC. Data deposition: PS. All authors edited and approved the final manuscript.

## Short Legends for Supporting Information

Table S1: HiFiasm Contig Assembly Statistics and Benchmarking Universal Single Copy Gene (BUSCO) Completeness in four *Macadamia* Species.

Table S2: Genome estimation statistics of four *Macadamia* species through K-mer analysis (using Jellyfish tool) and flow cytometry.

Table S3: Repeat Element Distribution across *Macadamia* Species

Table S4: Telomere distribution across all the four *macadamia* assemblies

Table S5: Distribution of Gene families (Fatty acid, cyanogenic and WRKY) across the four species of *Macadamia*.

Table S6: Distribution table of Orthologous gene clusters across the four *Macadamia* species and Telopea.

Figure S1 (a-d): K-mer profile (k = 21) spectrum analysis to estimate genome size of *M. jansenii, M. ternifolia, M. integrifolia and M. tetraphylla* generated from short read sequence data using Jellyfish and GenomeScope.

Figure S2: Dotplots illustrating the genomic comparison of *M. jansneii* Hi-C assembly (used as reference) against all the four assembled *Macadamia* genomes.

Figure S3: Dotplots illustrating the genomic comparisons between the haploid assemblies of each *Macadamia* species.

Figure S4: Species distribution graph of coding sequences of *M. jansenii*.

Figure S5: Multiple sequence aligmnet of Antimicrobial protein across the four *Macadamia* species. 01, 02, 03, 04,: represents AMP protein sequence *from M. jamsenii, M. ternifolia, M. integrifolia* and *M. tetraphylla,* respectively.

Figure S6: Distribution of unique and common orthologous gene clusters across the Macadamia species and Telopea.

Figure S7: Phylogenetic tree of *Macadamia* species with Telopea with number of orthogroups corresponding to each species

Figure S8: Self synteny of four *Macadamia* species, showing the collinearity of genes across the genome assemblies.

