## Supplemental material for "The genomes of the *Macadamia* genus"

|  | *M. jansenii* | | |  | *M. ternifolia* |  |  | *M. integrifolia* |  |  | *M. tetraphylla* |  |
| --- | --- | --- | --- | --- | --- | --- | --- | --- | --- | --- | --- | --- |
| Contig Assembly | | | | | | | | | | | | |
| Contig Assembly | Hap 1 | Hap2 | Collapsed | Hap 1 | Hap2 | Collapsed | Hap 1 | Hap2 | Collapsed | Hap 1 | Hap2 | Collapsed |
| N50 | 24 Mb | 14 Mb | 46 Mb | 27 Mb | 23 Mb | 47 Mb | 10 Mb | 9.8Mb | 45 Mb | 34 Mb | 27 Mb | 49 Mb |
| Length | 816 Mb | 776 Mb | 826 Mb | 810 Mb | 783 Mb | 827 Mb | 813 Mb | 797 Mb | 824 Mb | 821 Mb | 806 Mb | 839 Mb |
| Total Contigs | 879 | 363 | 779 | 883 | 361 | 803 | 1254 | 484 | 1049 | 809 | 262 | 742 |
| Contigs > 1Mb | 72 | 96 | 30 | 55 | 64 | 28 | 116 | 115 | 27 | 55 | 56 | 27 |
| Contigs >10Mb | 25 | 21 | 19 | 21 | 23 | 18 | 20 | 18 | 17 | 23 | 19 | 18 |
| BUSCO | 97.8% | 96.7% | 97.9% | 97.4% | 97.7% | 97.7% | 97.2% | 97.4% | 97.6% | 97.4% | 97.6% | 97.7% |
| Single BUSCO | 84.0% | 83.3% | 84.1% | 83.6% | 84.0% | 83.6% | 83.7% | 83.5% | 83.9% | 83.3% | 83.8% | 83.5% |
| Double busco | 13.8% | 13.4% | 13.8% | 13.8% | 13.7% | 14.1% | 13.5% | 13.9% | 13.7% | 14.1% | 13.8% | 14.2% |
| Fragmented | 0.6% | 0.7% | 0.7% | 0.8% | 0.9% | 0.8% | 0.9% | 0.7% | 0.6% | 0.8% | 0.7% | 0.7% |
| Missing | 1.6% | 2.6% | 1.4% | 1.8% | 1.4% | 1.5% | 1.9% | 1.9% | 1.8% | 1.8% | 1.7% | 1.6% |

Table S1: HiFiasm Contig Assembly Statistics and Benchmarking Universal Single Copy Gene (BUSCO) Completeness in four *Macadamia* Species

Table S2: Genome estimation statistics of four *Macadamia* species through K-mer analysis (using Jellyfish tool) and flow cytometry.

| **Methods to estimate genome size** | **Parameters** | ***M. jansenii*** | ***M. ternifolia*** | ***M. integrifolia*** | ***M. tetraphylla*** |
| --- | --- | --- | --- | --- | --- |
| **K-mer analysis** | Length | 643 Mb | 662 Mb | 676 Mb | 653 Mb |
|  | Heterozygous | 0.66% | 0.99% | 1.55% | 1.33% |
|  | Unique | 69.10% | 63.70% | 63.20% | 63.50% |
|  | Kcov | 16.3 | 7.74 | 9.33 | 6.76 |
|  | err | 0.35% | 0.14% | 2.27% | 0.15% |
|  | Dup | 0.65% | 0.26% | 0.65% | 0.49% |
| **Flow cytometry** | Length | 769 Mb | 775 Mb | 747 Mb | 796 Mb |
|  | Coefficient of variation | 1.19% | 0.49% | 0.80% | 0.62% |

Table S3: Repeat Element Distribution across *Macadamia* Species

|  | *M. jansenii* | | | *M. ternifolia* | | | *M. integrifolia* | | | *M. tetraphylla* | | |
| --- | --- | --- | --- | --- | --- | --- | --- | --- | --- | --- | --- | --- |
|  | Hap 1 | Hap2 | Collapsed | Hap 1 | Hap2 | Collapsed | Hap 1 | Hap2 | Collapsed | Hap 1 | Hap2 | Collapsed |
| SINE | 0.35% | 0.01% | 0% | 0.14% | 0.11% | 0.01% | 0.04% | 0.08% | 0.70% | 0.01% | 0.07% | 0.06% |
| LINE | 4.95% | 5.53% | 5.33% | 5.67% | 5.36% | 6.64% | 5.47% | 5.38% | 6.60% | 6.31% | 6.07% | 6.12% |
| LTR elements | 23.62% | 23.70% | 22.06% | 21.70% | 22.77% | 21.05% | 23.15% | 23.32% | 22.17% | 23.78% | 22.03% | 22.30% |
| DNA elements | 2.05% | 1.92% | 1.29% | 1.27% | 2.47% | 1.98% | 1.98% | 1.76% | 1.77% | 1.63% | 1.84% | 1.69% |
| Unclassified | 28.18% | 27.33% | 29.56% | 29.03% | 27.26% | 27.43% | 28.19% | 27.81% | 26.68% | 26.10% | 27.73% | 27.68% |
| Total interspersed repeats | 59.15% | 58.49% | 58.25% | 57.82% | 57.97% | 57.30% | 58.82% | 58.35% | 57.92% | 57.83% | 57.74% | 57.85% |
| Small RNA | 0.46% | 0.44% | 0.99% | 1.18% | 0.60% | 1.83% | 0.59% | 1.54% | 1.38% | 1.35% | 1.85% | 2.64% |
| Satellites | 0% | 0.02% | 0.03% | 0.01% | 0% | 0.01% | 0.02% | 0% | 0% | 0.02% | 0.01% | 0.03% |
| Simple repeats | 1.36% | 1.33% | 1.32% | 1.30% | 1.33% | 1.30% | 1.29% | 1.37% | 1.28% | 1.32% | 1.27% | 1.28% |
| Low complexity | 0.50% | 0.46% | 0.55% | 0.47% | 0.45% | 0.50% | 0.45% | 0.44% | 0.45% | 0.41% | 0.45% | 0.45% |
| Total Repeats | 61% | 61.00% | 61% | 61% | 61% | 61% | 61% | 61% | 61% | 61% | 61% | 62% |

Table S4: Telomere distribution across all the four *macadamia* assemblies

|  | M. jansenii | M. ternifolia | M. integrifolia | M. tetraphyllas |
| --- | --- | --- | --- | --- |
| Chr_01 | Telomere to telomere | Telomere to telomere | Telomere to telomere | Telomere to telomere |
| Chr_02 | Telomere to telomere | Telomere to telomere | Telomere to telomere | Telomere to NA |
| Chr_03 | Telomere to telomere | Telomere to telomere | Telomere to telomere | Na to telomere |
| Chr_04 | NA to telomere | NA to telomere | NA to Telomere | NA to telomere |
| Chr_05 | Telomere to telomere | Telomere to telomere | Telomere to telomere | Telomere to telomere |
| Chr_06 | Na to telomere | Telomere to telomere | Telomere to telomere | Telomere to telomere |
| Chr_07 | Telomere to Telomere | Telomere to telomere | NA to Telomere | Telomere to telomere |
| Chr_08 | Telomere to NA | Telomere to telomere | Telomere to telomere | Telomere to NA |
| Chr_09 | NA to telomere | Na to telomere | NA to Telomere | NA to telomere |
| Chr_10 | NA to telomere | Telomere to telomere | Telomere to NA | Telomere to NA |
| Chr_11 | Telomere to Telomere | Telomere to telomere | Telomere to telomere | Telomere to telomere |
| Chr_12 | Telomere to Telomere | Telomere to telomere | Telomere to telomere | Na to Telomere |
| Chr_13 | Telomere to Telomere | Telomere to telomere | Telomere to telomere | Telomere to telomere |
| Chr_14 | Telomere to Telomere | Telomere to NA | Telomere to telomere | Telomere to NA |

Table S5: Distribution of Gene families (Fatty acid, cyanogenic and WRKY) across the four species of *Macadamia*.

| (A) Fatty acid pathway genes | | | | |
| --- | --- | --- | --- | --- |
| Species | fatA | Fatb | SAD | KAS |
| *M. jansenii* | 9 | 7 | 18 | 10 |
| *M. ternifolia* | 8 | 8 | 23 | 10 |
| *M. integrifolia* | 10 | 11 | 28 | 0 |
| *M. tetraphylla* | 7 | 4 | 28 | 9 |

| (B) Cyanogenic pathway genes | | | | |
| --- | --- | --- | --- | --- |
| Species | CYP71 | CYP79 | BGLU | UGT |
| *M. jansenii* | 1 | 0 | 23 | 19 |
| *M. ternifolia* | 1 | 0 | 17 | 3 |
| *M. integrifolia* | 1 | 9 | 12 | 9 |
| *M. tetraphylla* | 1 | 12 | 16 | 0 |

| (C) WRKY gene |  |
| --- | --- |
| Species | Wrky gene |
| *M. jansenii* | 61 |
| *M. ternifolia* | 58 |
| *M. integrifolia* | 61 |
| *M. tetraphylla* | 59 |

Table S6: Distribution table of Orthologous gene clusters across the four *Macadamia* species and Telopea.

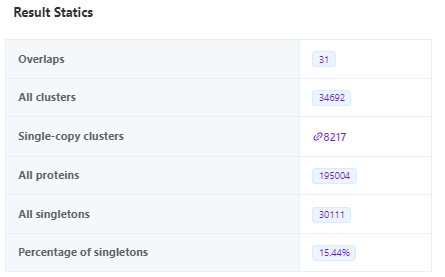

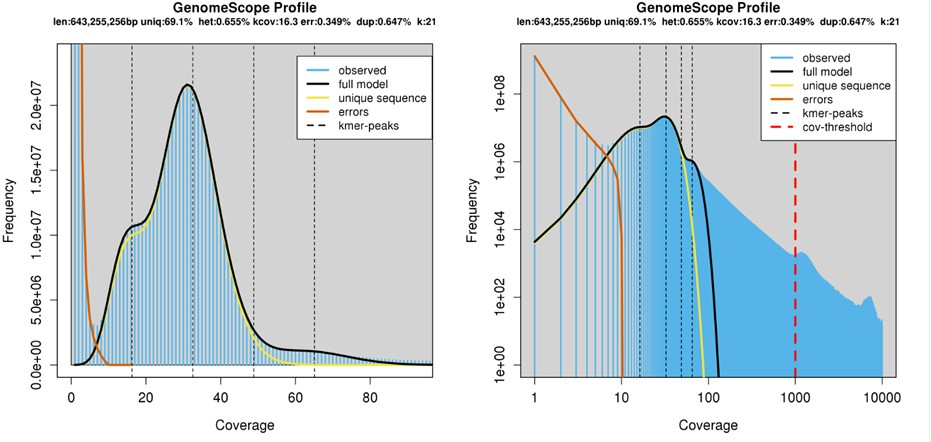

Figure S1 (a) : K-mer profile (k = 21) spectrum analysis to estimate genome size of *M. jansenii* generated from short read sequence data using Jellyfish and GenomeScope.

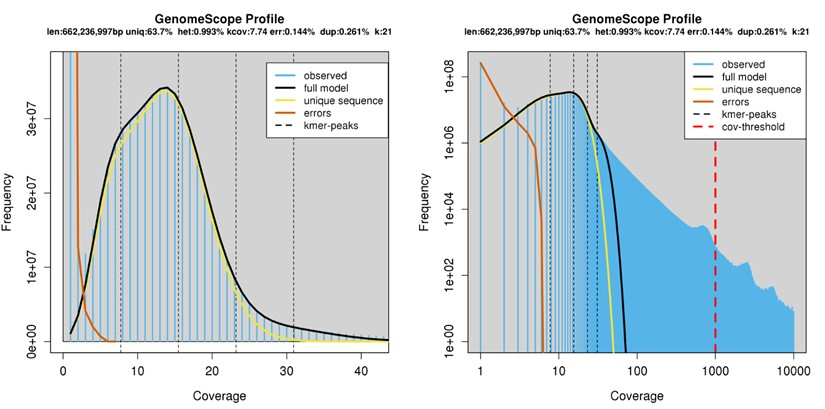

Figure S1 (b): K-mer profile (k = 21) spectrum analysis to estimate genome size of *M. ternifolia* generated from short read sequence data using Jellyfish and GenomeScope.

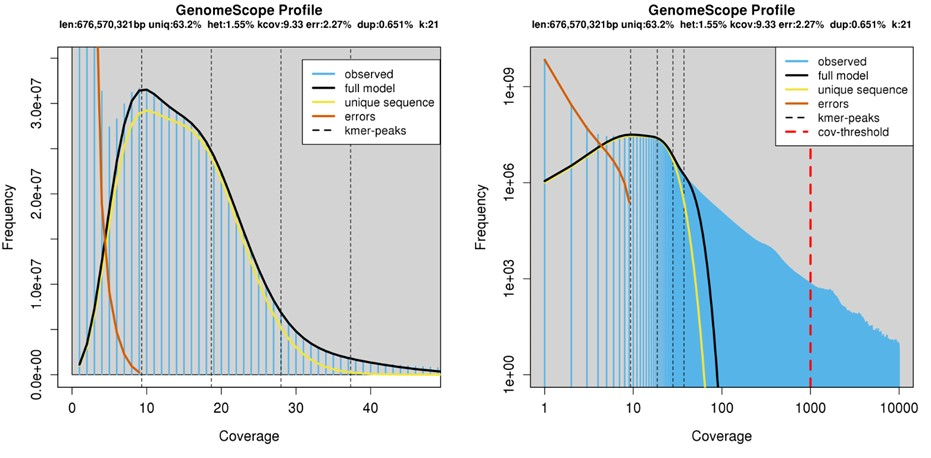

Figure S 1 (C): K-mer profile (k = 21) spectrum analysis to estimate genome size of *M. integrifolia* generated from short read sequence data using Jellyfish and GenomeScope.

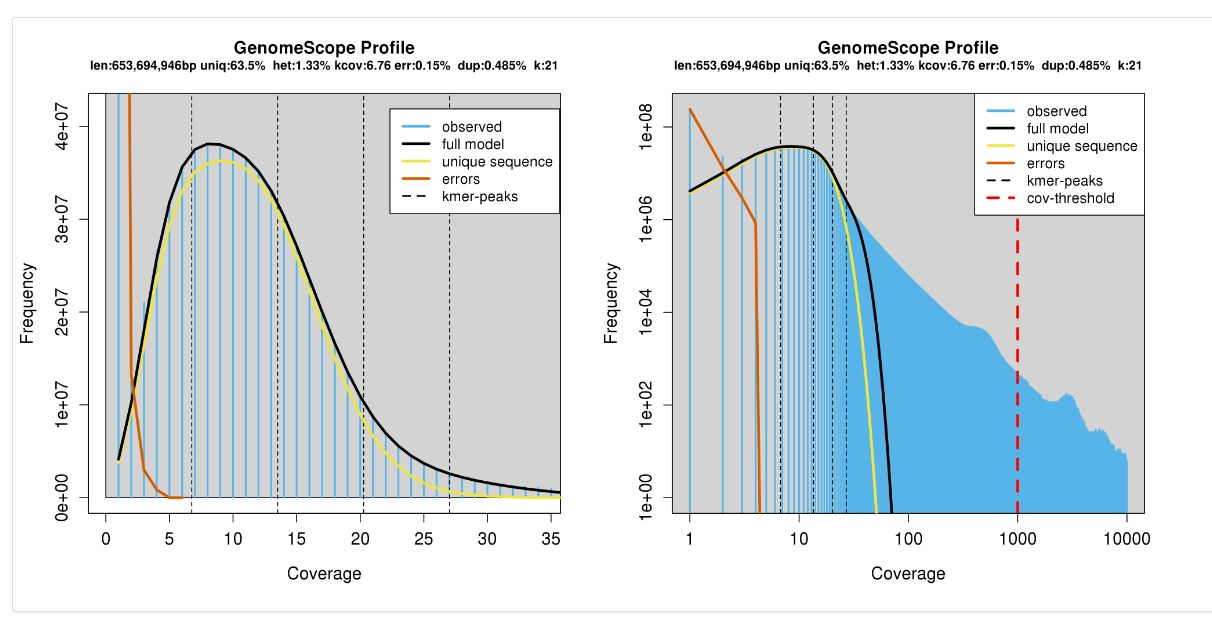

Figure S1 (d): K-mer profile (k = 21) spectrum analysis to estimate genome size of *M. tetraphylla* generated from short read sequence data using Jellyfish and GenomeScope.

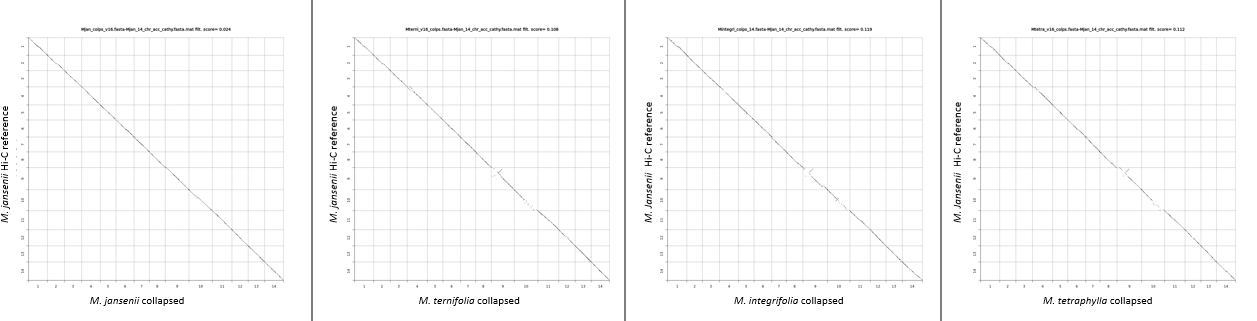

Figure S2: Dotplots illustrating the genomic comparison of *M. jansneii* Hi-C assembly (used as reference) against all the four assembled *Macadamia* genomes.

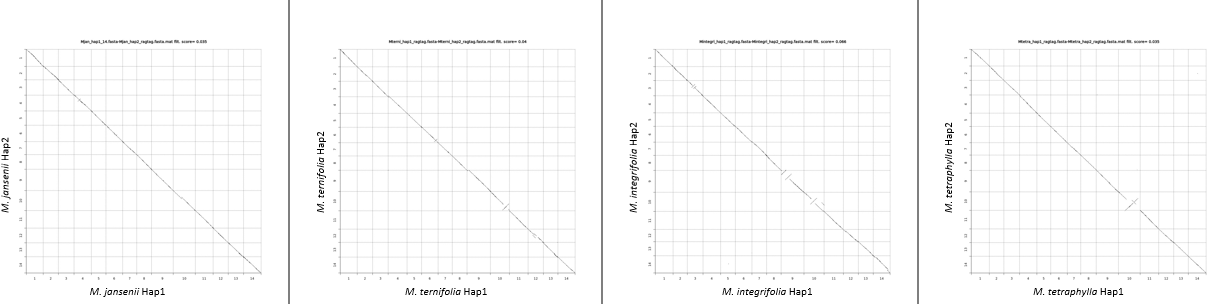

Figure S3: Dotplots illustrating the genomic comparisons between the haploid assemblies of each *Macadamia* species.

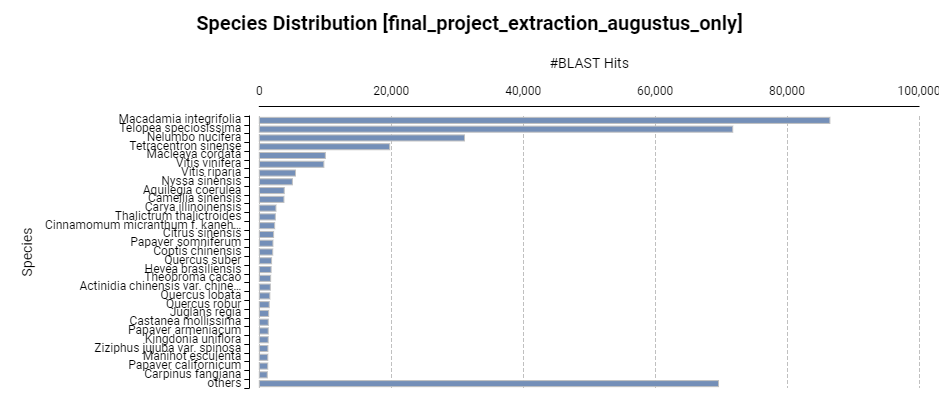

Figure S4: Species distribution graph of coding sequences of *M. jansenii*.

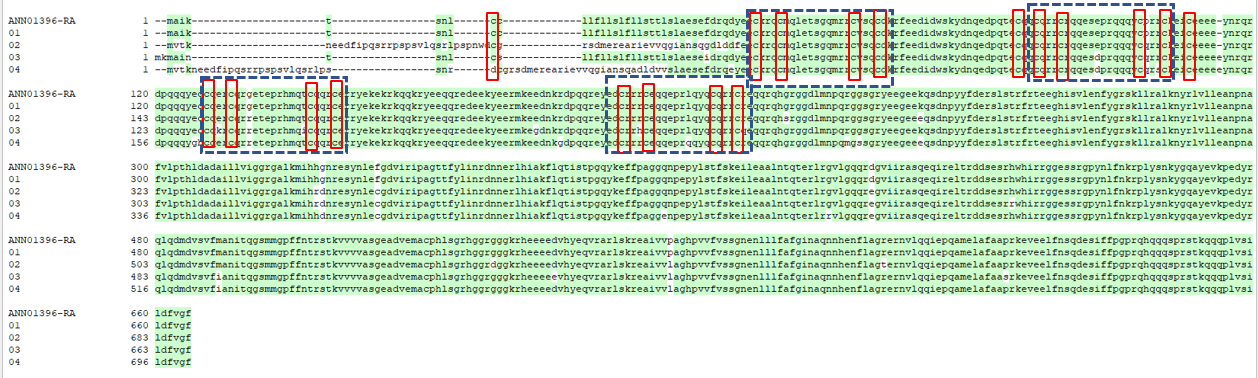

Figure S5: Multiple sequence aligmnet of Antimicrobial protein across the four *Macadamia* species. 01, 02, 03, 04, : represents AMP protein sequence from M. jamsenii, M. ternifolia, M. integrifolia and M. tetraphylla, respectively.

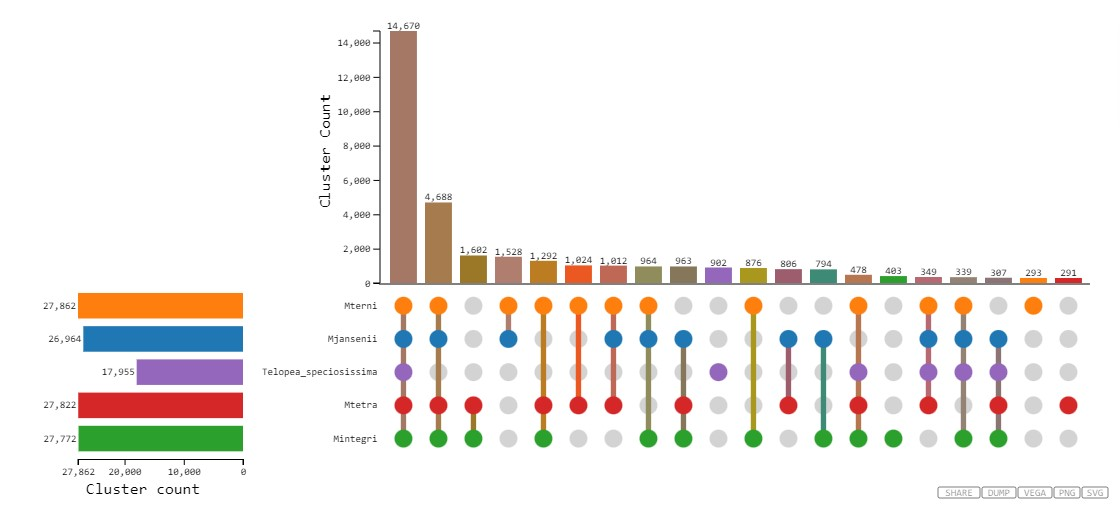

Figure S6: Distribution of unique and common orthologous gene clusters across the *Macadamia* species and *Telopea* .

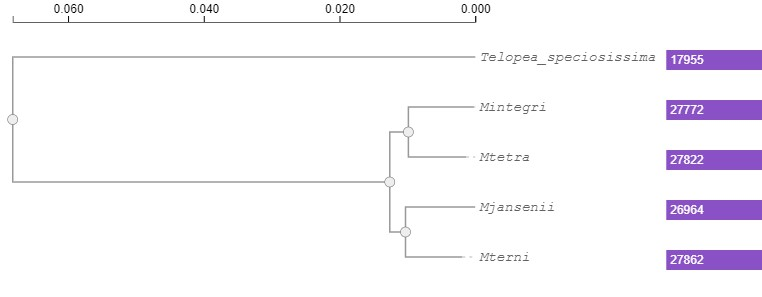

Figure S7: Phylogenetic tree of *Macadamia* species with Telopea, with number of orthogroups corresponding to each species in purple

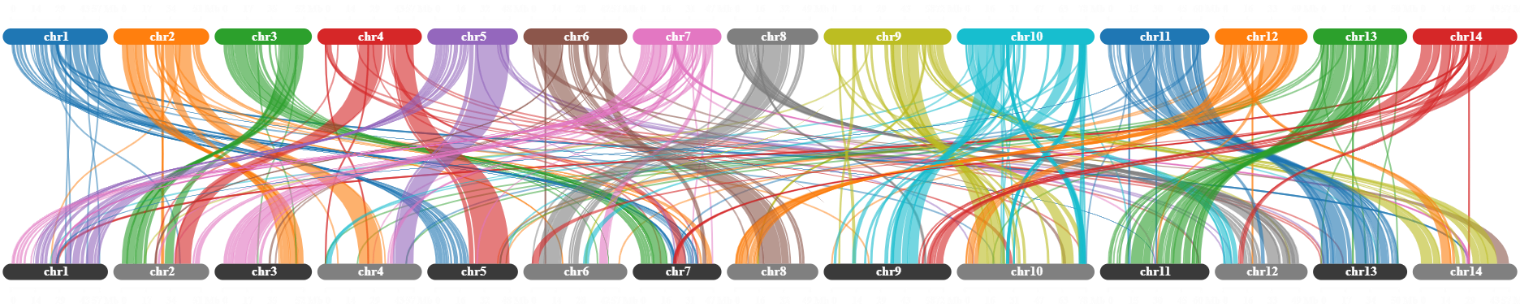

*M. jansenii*

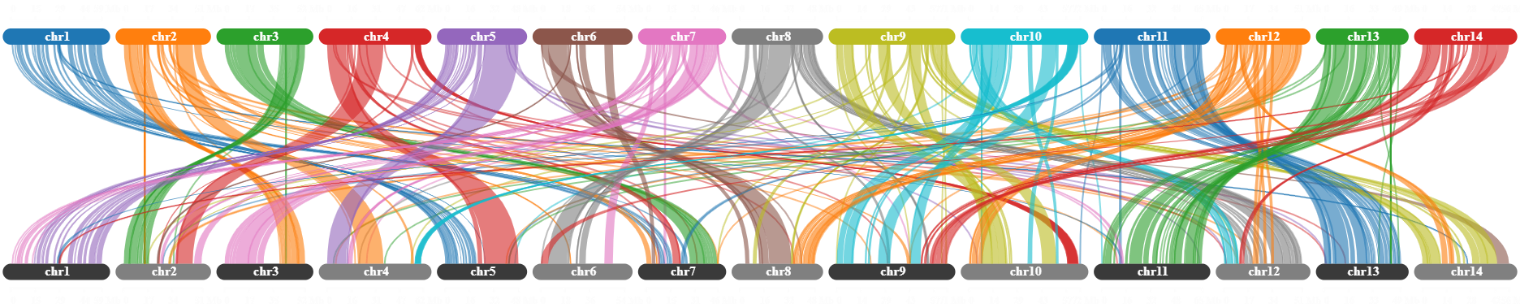

*M. ternifolia*

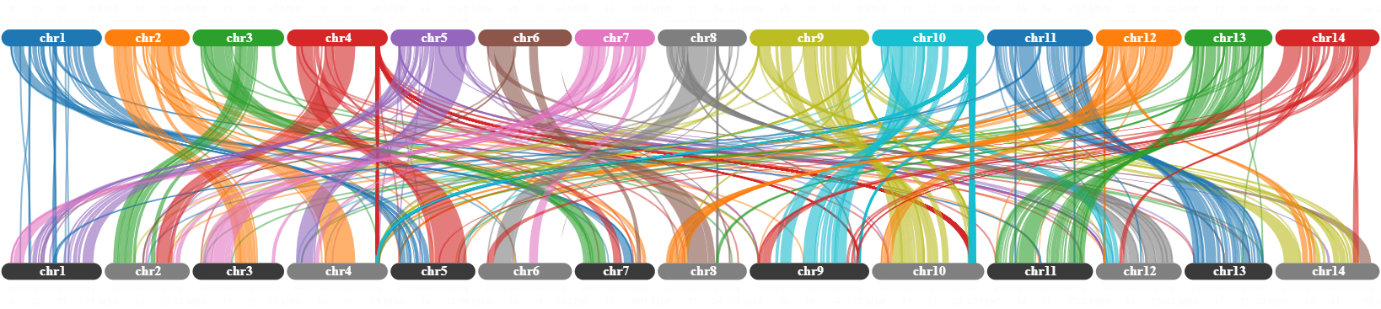

*M. tetraphylla*

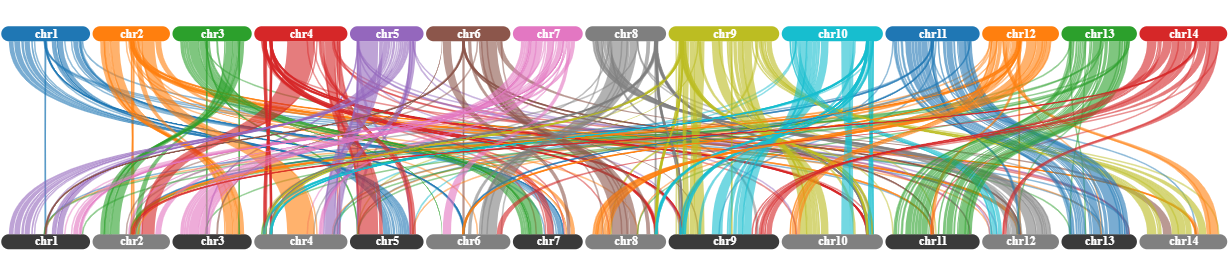

*M. integrifolia*

Figure S8: Self synteny of four *Macadamia* species, showing the collinearity of genes across the genome assemblies.
